# Structural basis for translational shutdown and immune evasion by the Nsp1 protein of SARS-CoV-2

**DOI:** 10.1101/2020.05.18.102467

**Authors:** Matthias Thoms, Robert Buschauer, Michael Ameismeier, Lennart Koepke, Timo Denk, Maximilian Hirschenberger, Hanna Kratzat, Manuel Hayn, Timur Mackens-Kiani, Jingdong Cheng, Christina M. Stürzel, Thomas Fröhlich, Otto Berninghausen, Thomas Becker, Frank Kirchhoff, Konstantin M.J. Sparrer, Roland Beckmann

**Affiliations:** Gene Center Munich, Department of Biochemistry, University of Munich, Munich, Germany; Institute of Molecular Virology, Ulm University Medical Center, Ulm, Germany; Laboratory of Functional Genome Analysis, University of Munich, Munich, Germany

## Abstract

SARS-CoV-2 is the causative agent of the current COVID-19 pandemic. A major virulence factor of SARS-CoVs is the nonstructural protein 1 (Nsp1) which suppresses host gene expression by ribosome association via an unknown mechanism. Here, we show that Nsp1 from SARS-CoV-2 binds to 40S and 80S ribosomes, resulting in shutdown of capped mRNA translation both *in vitro* and in cells. Structural analysis by cryo-electron microscopy (cryo-EM) of *in vitro* reconstituted Nsp1-40S and of native human Nsp1-ribosome complexes revealed that the Nsp1 C-terminus binds to and obstructs the mRNA entry tunnel. Thereby, Nsp1 effectively blocks RIG-I-dependent innate immune responses that would otherwise facilitate clearance of the infection. Thus, the structural characterization of the inhibitory mechanism of Nsp1 may aid structure-based drug design against SARS-CoV-2.

---

Coronaviruses (CoVs) are enveloped, single-stranded viruses with a positive-sense RNA genome, which infect a large variety of vertebrate animal species. Currently, seven CoV species from two genera (alpha and beta) are known human pathogens, four of which usually cause only mild respiratory diseases like common colds (*1-5*). Over the last two decades, however, three *Betacoronaviruses* (beta-CoVs) – the severe acute respiratory syndrome-coronavirus (SARS-CoV), the Middle East respiratory syndrome-coronavirus (MERS-CoV) and the novel severe acute respiratory syndrome–coronavirus 2 (SARS-CoV-2) – have emerged as the causative agents of epidemic and in the case of SARS-CoV-2 pandemic outbreaks of highly pathogenic respiratory diseases, which affect millions of people with a death toll amounting to hundreds of thousands worldwide (*6, 7*).

Coronavirus particles contain a single, 5’-capped and 3’-poly-adenylated RNA genome, which codes for two large overlapping open reading frames in gene 1 (ORF1a and ORF1b), as well as a variety of structural and nonstructural proteins at the 3’ end (*8, 9*). Following host infection, precursor proteins 1a and 1ab are translated and subsequently proteolytically cleaved into functional proteins, most of which play roles during viral replication (*10*). Amongst them is the N-terminal nonstructural protein 1 (Nsp1). Despite differences in protein size and mode of action, Nsp1 from alpha- and beta-CoVs display a similar biological function in suppressing host gene expression (*11-14*). SARS-CoV Nsp1 has been shown to induce a near-complete shutdown of host protein translation by a two-pronged strategy: first, it is able to bind the small ribosomal subunit and stall canonical mRNA translation at various stages during initiation (*15, 16*). Second, Nsp1 binding to the ribosome leads to template-dependent endonucleolytic cleavage and subsequent degradation of host mRNAs. Notably, shutdown of viral protein expression is circumvented via interactions between Nsp1 and a conserved region in the 5’ untranslated region (UTR) of viral mRNA through an unknown mechanism (*17*). Taken together, Nsp1 prevents all cellular anti-viral defense mechanisms that depend on the expression of host factors, including the interferon response. This drastic shutdown of the key parts of the innate immune system may facilitate efficient viral replication (*13, 18*) and immune evasion. Due to its central role in weakening the anti-viral immune response, SARS-CoV Nsp1 has been proposed as a potential therapeutic target (*19, 20*).

Here, we show how Nsp1 binds to the small ribosomal subunit and prevents canonical mRNA translation by blocking the mRNA entry tunnel. Moreover, we demonstrate that native targets of Nsp1 are non-translating, biogenesis-like or initiating 40S subunits as well as 80S ribosomes associated with CCDC124 and LYAR, trapped in their post-translocational state. We observed that expressing Nsp1 in cells thus results in an efficient shutdown of translation, in particular of innate immune response components. Our in-depth structural analysis may be useful for the design of drugs that target Nsp1 to achieve immune control and rapid elimination of invading Coronaviruses.

## Ribosome binding and translation inhibition of SARS-CoV-2 nsp1

About 84% aa sequence identity suggests that Nsp1 of SARS-CoV-2 may share similar properties and biological functions with its homolog from SARS-CoV (Fig. 1A). The C-terminal residues K164 and H165 in SARS-CoV are conserved in beta-CoVs and have been shown to be essential for 40S interaction since mutations to alanine abolish 40S binding and relieve translational inhibition (*16*). To confirm an analogous function of Nsp1 from SARS-CoV-2, we expressed and purified recombinant Nsp1 and the K164A/H165A mutant (Nsp1-mt) of both, SARS-CoV and SARS-CoV-2 from *E. coli*, and tested their binding efficiency to purified human ribosomal subunits (Fig. 1B). Nsp1 from both CoVs exhibited strong association to 40S subunits but not to 60S subunits, whereas both Nsp1-mt constructs failed entirely to bind (Fig. 1B and fig. S1A). Thus, ribosome binding to the 40S subunit is preserved and residues K164 and H165 of Nsp1 from both SARS-CoVs are important for this ribosome interaction. To further verify this, we expressed wildtype or mutant Nsp1 constructs in human HEK293T cells and analyzed ribosome association by sucrose-gradient centrifugation. Consistent with the behavior *in vitro*, Nsp1 of CoV and CoV-2 co-migrated with 40S ribosomal subunits and 80S ribosomes, but not with actively translating polyribosomes. In contrast, the mutant constructs barely penetrated the gradient, indicative of loss of their affinity for ribosomes (Fig. 1C). Notably, compared to the control the polysome profiles showed a shift from translating polyribosomes to 80S monosomes in presence of Nsp1, indicative of global inhibition of translation. This effect was less pronounced for the two Nsp1-mt constructs. Next, we performed *in vitro* translation assays of capped reporter mRNA in cell free translation extracts from human cells (HeLa S3) or rabbit reticulocytes in the presence of Nsp1 or Nsp1-mt. Probing for the translation products by Western blotting revealed a complete inhibition of translation by Nsp1 and only weak effects in presence of Nsp1-mt constructs (Fig. 1D and fig. S1B).

**Fig. 1.**
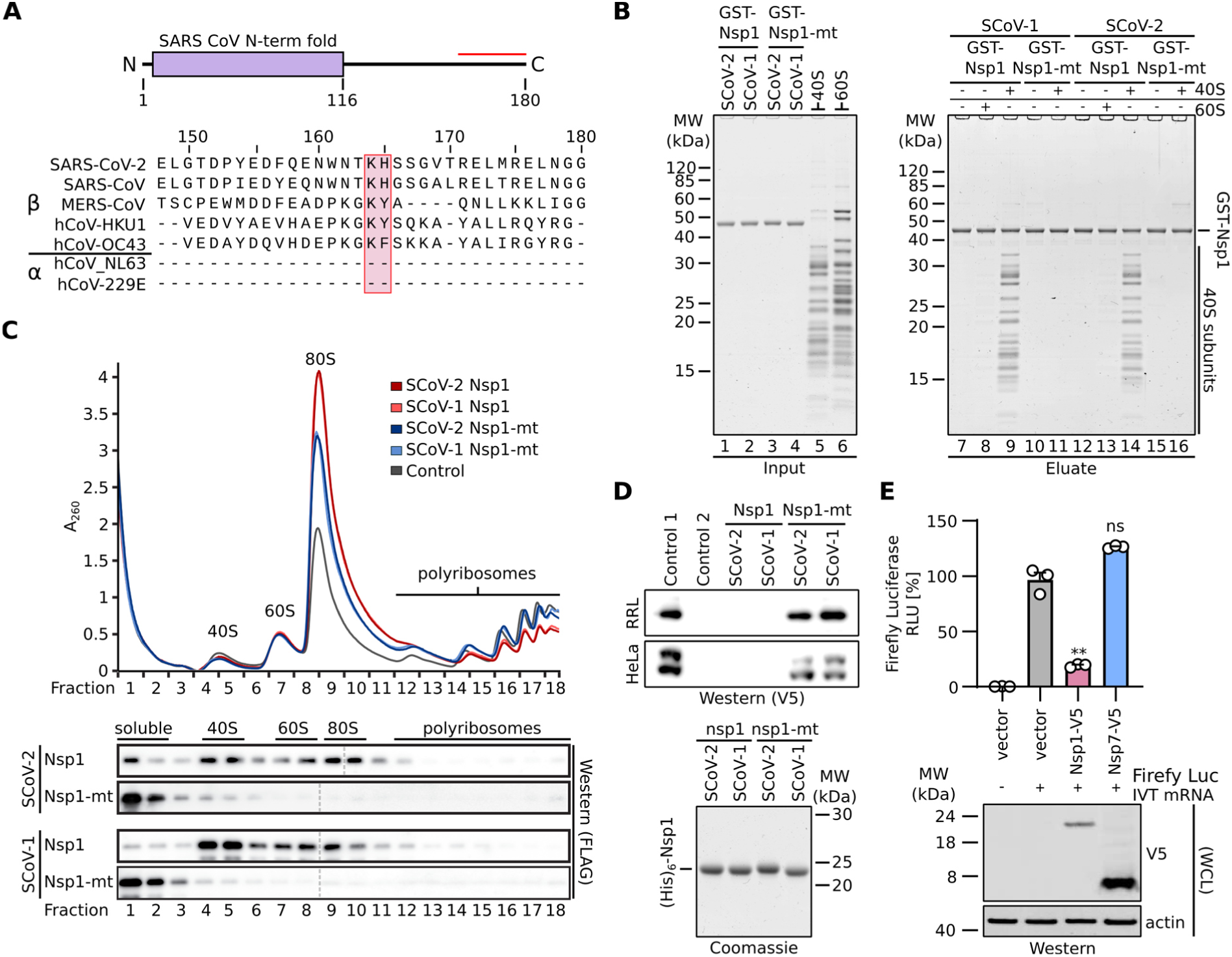
Nsp1 interacts with 40S ribosomal subunits and inhibits translation. (**A**) Domain organization of Nsp1 and sequence alignment of the C-terminal segment (red line) of Nsp1 from seven human pathogenic CoVs. The KH motif is marked with a red box. (**B**) Nsp1 interacts with 40S ribosomal subunits. *In vitro* binding assay of N-terminal GST-TEV (GST) tagged Nsp1 and Nsp1-mt from SARS-CoV (SCoV-1) and SARS-CoV-2 (SCoV-2) with human 40S and 60S ribosomal subunits. Coomassie stained SDS-PAGE of the inputs (lane 1-6) and eluates (lane 7-16) are shown. (**C**) Polyribosome gradient analysis of HEK293T cell lysate (Control) and lysate from HEK293T cells transiently transfected with N-terminal 3xFLAG-3C tagged Nsp1 and Nsp1-mt constructs from SCoV-1 and SCoV-2. Western blot analysis of the indicated gradient fractions stained with anti-FLAG antibody are shown below and separate blots are indicated with dashed lines. Fractions containing 40S, 60S, 80S and polyribosomes are labeled. (**D**) Cell-free *in vitro* translation of a capped reporter mRNA with rabbit reticulocytes (RRL) and HeLa S3 lysate. The translation product was analyzed by western blot analysis using a anti-V5 antibody (upper panel). Control 1 and Control 2, with and without capped reporter mRNA, respectively. Coomassie stained SDS-PAGE of the applied N-terminal (His)_6_-TEV (His_6_) tagged Nsp1 constructs are shown below. (**E**) Luciferase reporter gene assay of HEK293T cells transiently transfected with indicated plasmids (empty vector, Nsp1-V5, Nsp7-V5) and *in vitro* transcribed Firefly luciferase mRNA. Bars represent the mean of n=3±SEM. RLU, relative light units, normalized to empty vector set to 100% (top panel). Western blot of whole cell lysates (WCL) stained with anti-V5 and anti-actin antibodies (bottom panel). Unpaired student’s t-test (Welch correction), ns, non significant; **, p<0.001

To test the inhibitory effect of Nsp1 on translation in cells, we expressed Nsp1 and as a control Nsp7, which is also derived from SARS-CoV-2 Orf1a/b, in HEK293T cells and monitored translation of a co-transfected capped luciferase reporter mRNA. Consistent with the results of the *in vitro* assays, we observed a near complete loss of translation in presence of Nsp1 (Fig. 1E). This phenotype was confirmed for codon-optimized, non-codon-optimized Nsp1, as well as differently tagged versions of Nsp1 (fig. S1, C and D). In summary, Nsp1 from both, SARS-CoV and SARS-CoV-2 binds 40S and 80S ribosomes and disrupt cap-dependent translation. Moreover, the conserved KH motif close to the C-terminus of Nsp1 is crucial for ribosome binding and translation inhibition.

## In vitro reconstituted and native nsp1-40S and - 80S ribosome complexes

To elucidate the molecular interaction of SARS-CoV-2 Nsp1 with human ribosomes, we reconstituted a complex from purified, recombinant Nsp1 together with purified human 40S ribosomal subunits and determined its structure by cryo-EM at an average resolution of 2.6 Å (Fig. 2, A and B and figs. S2 and S3). In addition to the 40S ribosomal subunit, we observed density corresponding to two α-helices inside the ribosomal mRNA entry channel, which could be unambiguously identified as the C-terminal part of Nsp1 from SARS-CoV-2 (Fig. 2C).

**Fig. 2.**
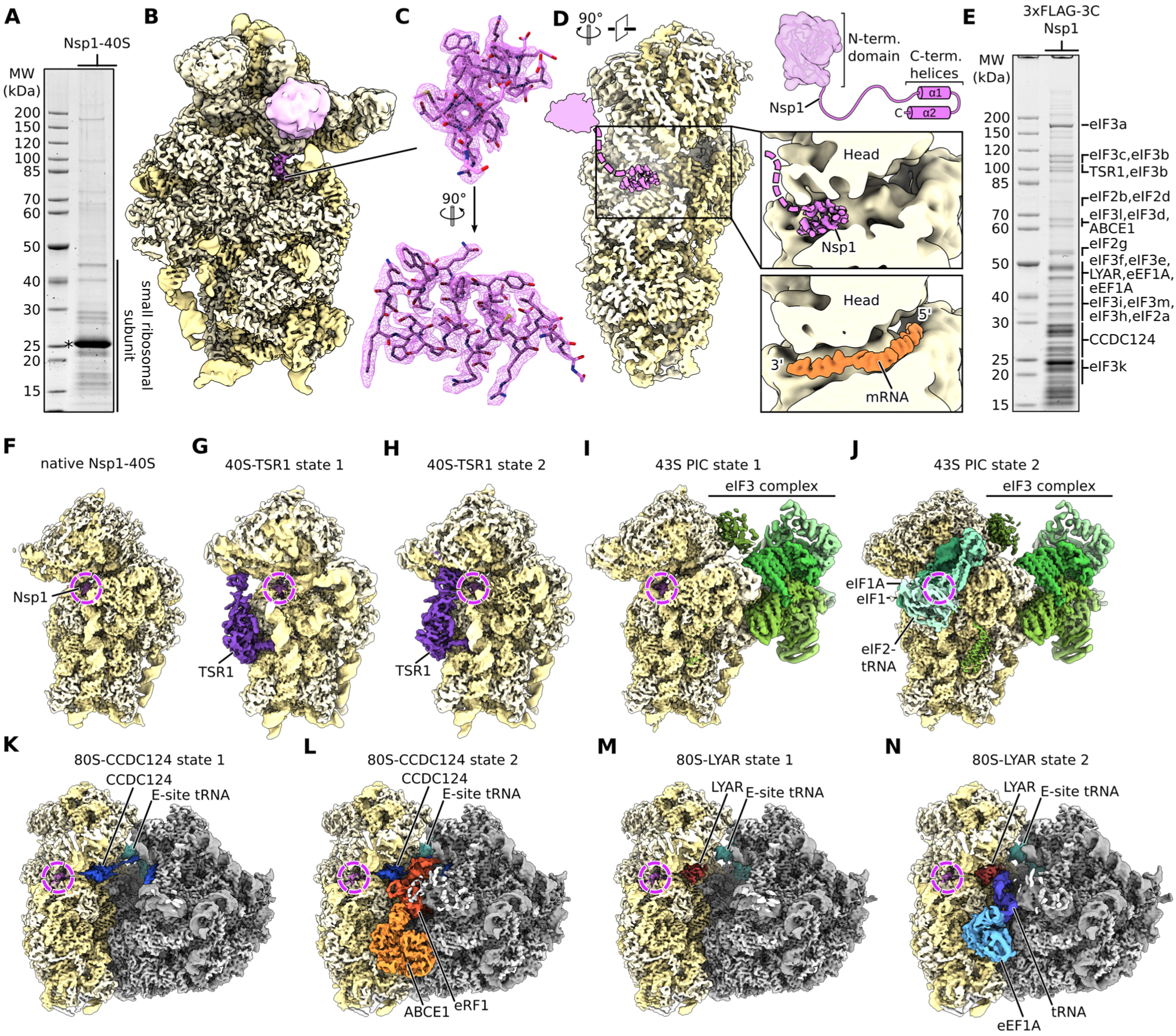
Cryo-EM structures of nsp1-bound ribosomal complexes. (**A**) SDS-PAGE analysis of reconstituted Nsp1-40S complexes, Nsp1 is labeled with an asterisk. (**B**) Reconstituted Nsp1-40S structure with Nsp1 in pink and ribosomal RNA and proteins in yellow. Additional density between uS3 and h16 assigned to the N-terminal fold of Nsp1. (**C**) C-terminal helix 1 and 2 of Nsp1 with corresponding density. (**D**) Cross-section of the 40S highlighting the central position of Nsp1 within the mRNA tunnel. The putative position of the N-terminal domain of Nsp1 is schematically indicated (models based on PDB-2HSX and PDB-6Y0G) (**E**) SDS-PAGE analysis of affinity purified Nsp1-ribosomal complexes. Proteins identified in the cryo-EM structures were labeled according to mass spectrometry analysis. (**F** to **N**) Cryo-EM maps of affinity purified Nsp1-ribosomal complexes. Additional factors are colored and labeled accordingly.

In proximity to the helical density, we observed undefined globular density between rRNA helix h16 and ribosomal proteins uS3 and uS10. The dimensions of this extra density roughly matched the putative dimensions of the globular N-terminal domain of Nsp1 (Fig. 2, C and D), based on a structure of the highly similar N-terminus of Nsp1 from SARS-CoV previously determined by NMR (*21*). However, the resolution of this region in our cryo-EM density map was insufficient for unambiguous identification. The C-terminus of Nsp1 is located close to the so called “latch” between the rRNA helix h18 of the body and h34 of the head of the 40S subunit, which influences mRNA accommodation and movement during translation initiation (*22, 23*). When bound at this position, the Nsp1 C-terminus blocks regular mRNA accommodation, thus providing an explanation of Nsp1’s activity in host translation shutdown (Fig. 2D).

To characterize the ribosomal targets and the mode of interaction of Nsp1 in the human cellular context, we expressed N-terminally 3xFLAG tagged Nsp1 in HEK293T cells and affinity purified associated native complexes for analysis by cryo-EM and mass spectrometry (Fig. 2E, figs. S2 and S3 and Data S1). Structural analysis revealed 40S and 80S ribosomal complexes in nine compositionally different states (Fig. 2, F to N). Importantly, all of them displayed density for the Nsp1 C-terminus in the same position and conformation as observed in the *in vitro* assembled complex and lacked density corresponding to mRNA. Yet, apart from these communalities the native complexes were of a diverse nature and, besides the 43S pre-initiation complexes, rather unexpected.

The Nsp1-bound 40S ribosomal complexes could be divided into three major populations. The first represented idle Nsp1-40S complexes (Fig. 2F), essentially resembling the *in vitro* reconstituted complex. The second population were unusual, pre-40S-like complexes (Fig. 2, G and H), which contained the cytosolic ribosome biogenesis factor TSR1 bound in two distinct conformations between the 40S head and body (*24, 25*). Notably, these complexes did not resemble any known *bona-fide* biogenesis intermediates. The third population represented 43S pre-initiation complexes (PICs), which could be further divided into eIF3-containing PICs and complexes containing in addition to eIF3 also eIF1A, eIF1, and a fully assembled ternary eIF2-tRNA_i_-GTP complex (Fig. 2, I and J) (*26-28*). This stable association of Nsp1 in the cell with multiple different intermediates states of translation initiation besides empty 40S ribosomal complexes is in agreement with the proposed role of Nsp1 as an inhibitor of translation initiation (*15*).

The Nsp1-bound 80S complexes could be divided into two major populations of translationally inactive ribosomes, which have to our knowledge not been observed before. The first population (Fig. 2, K and L and fig. S4, A to E) contained the protein coiled-coil domain containing short open reading frame 124 (CCDC124), a homolog of the ribosome protection and translation recovery factor Lso2 in *Saccharomyces cerevisiae* (*29*). A similar complex of inactive 80S ribosomes bound to CCDC124 was recently described (*30*). In contrast to the known hibernation complex, however, a subpopulation of the Nsp1-bound complex contained in addition the ribosome recycling factor and ABC-type ATPase ABCE1 (*31-33*) together with class I translation termination factor eRF1 in an unusual conformation (fig. S4, C to E). In addition, the previously unresolved, flexible C-terminal part of CCDC124 was stably bound to the ribosomal A-site in this complex. Together, this complex might represent a novel ribosome recycling-like state.

The second major population of Nsp1-bound 80S ribosomes (Fig. 2, M and N) contained the cell growth regulating nucleolar protein LYAR, which was previously implicated in processing of pre-rRNA and in negative regulation of antiviral innate immune response (*34, 35*). We found the C-terminus of LYAR occupying the ribosomal A-site, similar to CCDC124 (Fig. 2, M and fig. S4, F and G). Furthermore, we identified a subpopulation among the LYAR-bound inactive 80S ribosomes that contained a ternary eIF1A-GTP-tRNA complex (Fig. 2, N and fig. S4, H to K). This ternary complex was in an unusual conformation, with the anticodon loop contacting an α-helix of the LYAR C-terminus. Such a complex has not been described yet and its functional relevance is unknown.

Taken together, we found Nsp1 bound to the mRNA entry channel of translationally inactive 80S ribosomes. Among them were highly unusual complexes, for which it is unclear, whether they were a result of the presence of Nsp1, or whether they occur naturally and have an increased affinity for Nsp1 due to their distinct conformation or lack of mRNA.

## Molecular basis of nsp1-ribosome interaction and inhibition

We observed the same binding mode of Nsp1 to the 40S subunit in all ribosomal complexes. The C-terminal domain of Nsp1 (Nsp1-C) was rigidly bound inside the mRNA entry channel, which, at this location, is constituted by the rRNA helix h18 and the ribosomal proteins uS5 of the 40S body and uS3 of the 40S head. The local resolution of 2.6 Å (fig. S3) allowed for detailed analysis of the molecular interactions of Nsp1 with the ribosome (Fig. 3A).

**Fig. 3.**
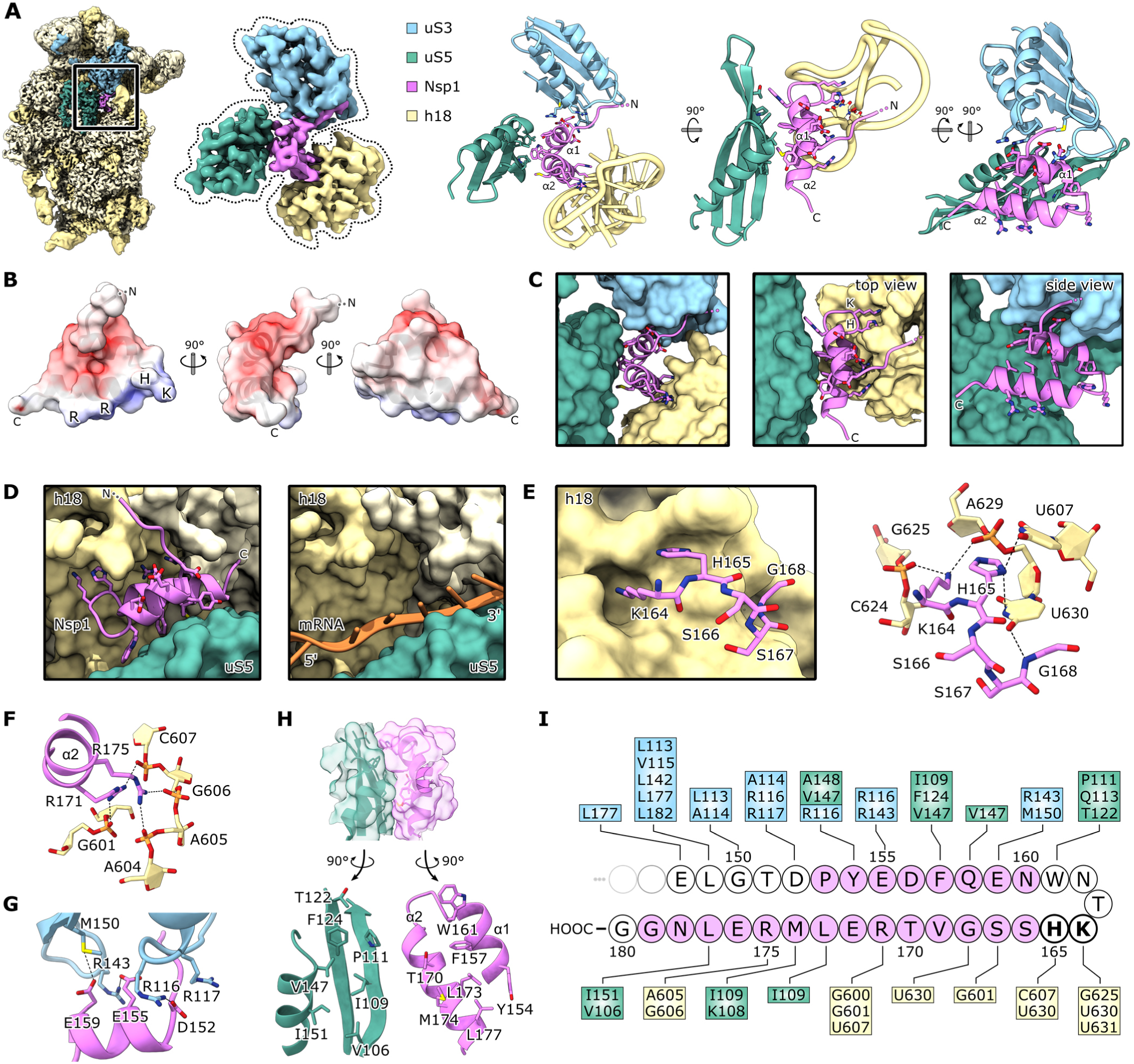
Molecular basis of nsp1 ribosome interaction and inhibition. (**A**) Cryo-EM map of *in vitro* reconstituted Nsp1-40S and segmented density of Nsp1-C, uS3 (97-153,168-189), uS5 (102-164) and rRNA helix h18 with the corresponding models; interacting residues are shown as sticks. (**B**) Nsp1-C surface, colored by electrostatic potential from −5 (red) to +5 (blue). (**C**) Model of Nsp1-C and surface representation of the models of uS3 (97-153,168-189), uS5 (102-164) and rRNA helix h18. Molecular interactions between Nsp1 and the ribosome. (**D**) mRNA entry channel, 40S head is removed. Nsp1-C occupies the mRNA path. (**E**) K164 and H165 of Nsp1 bind to a pocket on h18. (**F**) R171 and R175 of Nsp1 bind to the phosphate backbone of h18. (**G**) Negatively charged residues D152, E155 and E159 of α1 interact with uS3. (**H**) The hydrophobic interface of α1 and α2 binds to a hydrophobic patch on uS5. (**I**) Schematic summary of the interaction of Nsp1-C with uS3, uS5 and h18; residues belonging to α1 and α2 are colored in pink.

The shorter, first α-helix of Nsp1-C (α1; residues 154-160) interacted with uS3 and uS5. The helix was followed by a short loop, which contained the essential KH motif that interacted with h18. Notably, this part of h18 belongs to the so-called “530-loop”, which is actively participating in ribosomal decoding, and has been reported to resemble a conserved structural motif in the 3’-UTR of beta-CoVs (*36*). The second, larger α-helix of Nsp1-C (α2; residues 166-179) also interacted with rRNA h18 and connected back to uS5 at its C-terminal end. The two helices stabilized each other through hydrophobic interactions. The electrostatic potential on the Nsp1-C surface displayed three major patches (Fig. 3B). A negatively charged patch on α1 facing positively charged residues on uS3, a positively charged patch on α2 facing the phosphate backbone of h18 and a hydrophobic patch at the α1-α2 interface, which was exposed to hydrophobic residues on uS5. In addition to the matching surface charge, also the shape of Nsp1-C perfectly matched the shape of the mRNA channel and thus completely overlapped with the regular mRNA path (Fig. 3, C and D). Together, this explains the strong inhibitory effect on translation observed *in vitro* and *in vivo*. A key interaction was established through the KH motif, which was bound to a distinct site on rRNA helix h18 (Fig. 3, C and E); K164 of Nsp1 inserted into a negatively charged pocket, constituted mainly by the phosphate backbone of rRNA bases G625 and U630, whereas H165 stacked in between U607 and U630. The base U630 was stabilized in this position through interaction with the backbone of G168 of Nsp1. The strict dependence of the ribosome interaction of Nsp1 and its inhibitory function on the KH motif suggests this pocket as a potential drug target. Further interactions to h18 were established through R171 and R175 of Nsp1, which formed salt bridges to the backbone phosphates of G601, C607, A605 and G606 of h18 (Fig. 3F). The interactions of Nsp1-C and uS3 were established through salt bridges and hydrogen bonds between D152, E155, E159 of Nsp1 and R116, R143 and M150 of uS3 (Fig. 3G). The interactions of Nsp1-C with uS5 were established along a hydrophobic surface of ∼440 Å^2^ involving residues Y154, F157, W161, T170, L173, M174, L177 of Nsp1 and residues V106, I109, P111, T122, F124, V147, I151 of uS5 (Fig. 3H). Taken together, Nsp1 established a multitude of specific molecular contacts (summarized in Fig. 3I) to rigidly anchor into and thereby obstruct the mRNA entry channel. Therefore, disruption of one or multiple of the interfaces through competition with a peptide or small molecule may represent a potential therapeutic strategy against multiple beta-CoVs.

## SARS-CoV-2 Nsp1 efficiently shuts down the type-I interferon innate immune response

Type-I interferon induction and signaling represents one of the major innate anti-viral defense pathways, ultimately leading to the induction of more than 600 interferon-stimulated genes (ISGs) (*37*). The type I interferon pathway is activated by the detection of CoVs via the innate immune sensor RIG-I (*37-39*). To assess the effects of SARS-CoV-2 Nsp-1 on type-I interferon induction, we stimulated HEK293T cells with Sendai Virus (SeV), a well-known trigger of RIG-I-dependent signaling (*37, 40*). Expression of Nsp1 completely inhibited the SeV triggered induction of the human interferon-beta (IFN-β) promotor, while Nsp7 had no significant effect (Fig. 4A). Measles virus (MeV) V protein, which inhibits immune sensors like MDA5 (*41*), but has little effect on RIG-I-dependent signaling, was included as control for specific induction of RIG-I-dependent signaling. Notably, the presences of Nsp1 did not impact the induction of the endogenous IFN-β or Interleukin 8 (IL-8) mRNA upon SeV stimulation. In contrast the release of IFN-β and IL-8 (Fig. 4, B and C) was severely impaired upon expression of Nsp1 (Fig. 4, B and C). Again, Nsp7 served as a negative control. IFN-β-mediated induction of expression from the interferon stimulated response element (ISRE), which is part of the promoter of most ISGs, was effectively shut down by Nsp1 and MeV V (Fig. 4D) (*42*). These results support that SARS-CoV-2 Nsp1 blocks not only primary induction of the cytokines, but also expression of ISGs upon exogenous cytokine stimulation by specific disruption of the translation step. Notably, not all innate immune responses require active translation for function. Among these responses is autophagy, where all components required for anti-viral activity are already present in the cytoplasm and activated by post-translational modifications. Consequently, autophagy is barely affected by the expression of Nsp1 (fig. S5). Tripartite Motif Protein 32 (TRIM32), which was used as a positive control, induced a strong autophagic response (*43*), whereas the negative control Nsp7 had no effect.

**Figure 4:**
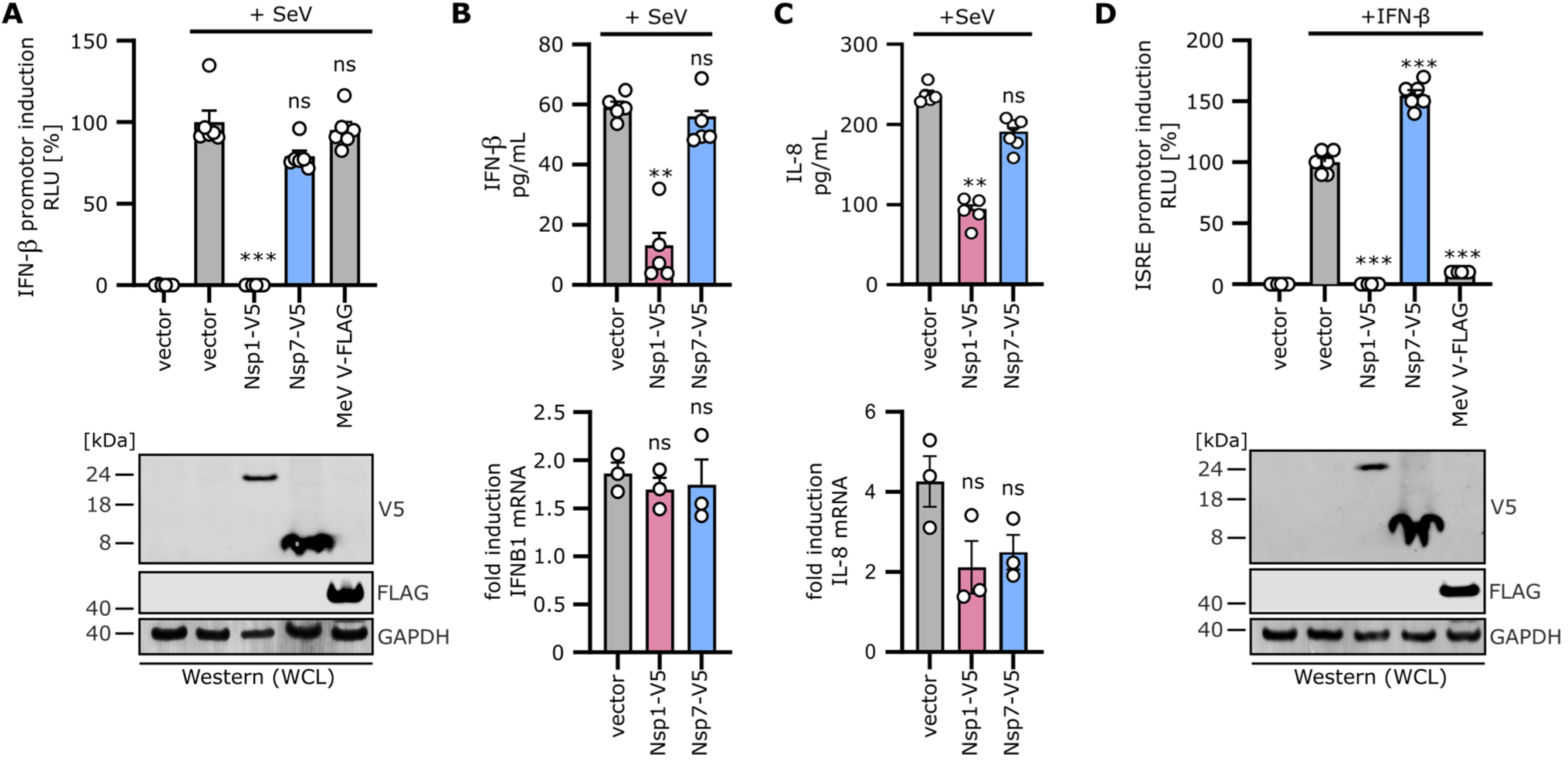
Inhibition of the innate immune response by SARS-CoV-2 nsp1. (**A**) IFN-β-promotor controlled Luciferase reporter gene assay. HEK293T cells were transiently transfected with indicated plasmids and 24 h post transfection cells were either infected with Sendai Virus (SeV) or left uninfected. Firefly Luciferase was quantified 28 h post infection. Bars represent the mean of n=6±SEM. RLU, relative light units, stimulated vector control set to 100% (top panel). Immunoblot of whole cell lysates stained with anti-V5, anti-FLAG and anti-GAPDH antibodies (bottom panel). (**B** and **C**) ELISA of IFN-β (B) or IL-8 (C) released by HEK293T cells transiently transfected with indicated vectors and infected with SeVs (top panel) for 24 h. Bars represent the mean of n=5±SEM. qPCR of the respective mRNA induction (bottom panel). Bars represent the mean of n=3±SEM. (**D**) Interferon Stimulated Response Element (ISRE)-promotor controlled Luciferase reporter assay. HEK293T cells were transiently transfected with indicated plasmids and treated 24h post transfection with 1000 U/mL IFN-β or left untreated. Firefly Luciferase expression was quantified 8 h post stimulation. Bars represent the mean of n=6±SEM, the stimulated vector control set to 100% (top panel). Immunoblot of whole cell lysates stained with anti-V5, anti-FLAG and anti-GAPDH antibodies (bottom panel).

Taken together, these results demonstrate that the type-I interferon system and consequently anti-viral ISG expression is almost completely shut down by SARS-CoV-2 Nsp1 at the level of translation. Thus, our data establish that SARS-CoV-2 uses Nsp1 as a major inhibitor of anti-viral innate immune responses.

## Discussion

Our data establish that one of the major immune evasion factors of SARS-CoV-2, Nsp1, efficiently interferes with the cellular translation machinery in order to shut-down the production of host proteins including antiviral defense factors. As the molecular basis we discovered the obstruction of the mRNA entry channel by the C-terminal domain of Nsp1, which employs a highly specific interaction with uS3, uS5 and rRNA helix 18 on the 40S small ribosomal subunit. Here, the KH motif establishes a key interaction with a distinct pocket on h18, which explains why it is essential for Nsp1 function.

Sensing of Coronaviruses by RIG-I mounts an innate immune response that - if not blocked by the virus - would efficiently restrict viral replication (*44*). A central target of the translational shutdown is the innate immune response, in particular the RIG-I-dependent type-I interferon signaling cascade, which depends on the induction of genes and their subsequent translation to exert anti-viral action. Although SARS-CoV-2 encodes additional potential inhibitors of the innate immune system, a loss of Nsp1 function might thus enable the host cell to efficiently restrict viral replication and consequently prevent the development of a severe respiratory disease. Recombinant SARS-CoV-2 bearing mutations in Nsp1 could be used to clarify the relative contribution of Nsp1 to immune evasion in the viral context. In combination with animal models, this would also allow an assessment of the importance of Nsp1 on viral replication and pathogenesis *in vivo*.

Because the inhibition of translation is strictly dependent on the ribosome association of Nsp1, interfering with this interaction could represent a powerful approach to fight COVID-19. To this end, our data at 2.6 Å resolution provide a starting point for rational structure-based drug design, in particular considering the binding pocket of the critical KH motif on rRNA helix h18. Ribosomes are established targets of a plethora of small molecules such as antibiotics (*45*), and to the best of our knowledge, the pocket on h18 is not used by any host factor during normal translation or ribosome biogenesis. Thus, a small molecule masking this Nsp1 binding pocket, may prevent association of Nsp1 with the ribosome, but still allow for normal cellular translation.

Still, important questions remain: is translation of all cellular mRNAs equally affected by Nsp1? Is Nsp1 of SARS-CoV-2 also causing modification and cleavage of cellular mRNAs as suggested for the SARS-CoV homolog (*16*)? Most interestingly, however, how does the virus ensure that the production of its own, viral proteins is still possible in the presence of Nsp1? Research on SARS-CoV hints at a role of the 5’ untranslated region, which is also present on all SARS-CoV-2 mRNAs, in circumventing the ribosome blockage by Nsp1 (*46*). Addressing these questions will be important for understanding SARS-CoV-2 infection and a better knowledge of the molecular interactions of this viral pathogen with the host cell might allow for the development of safe and effective approaches to combat it.

## Acknowledgments

Sendai virus was kindly provided by Georg Kochs and Daniel Sauter. Luciferase Reporter constructs were provided by Karl-Klaus Conzelmann. We thank Susanne Engelhart, Kerstin Regensburger, Martha Meyer, Regina Burger, Nicole Schrott, Daniela Krnavek, Miwako Kösters, Charlotte Ungewickell and Susanne Rieder for excellent technical assistance.

## Author contributions

R.Be., K.S., M.T., R.Bu. and M.A. designed the study; O.B. collected cryo-EM data, M.T., R.Bu. and M.A. prepared cryo-EM samples and processed cryo-EM data; R.Bu., M.A. and J.C. built molecular models; T.D. performed *in vitro* translation assays with help from H.K.; M.T. generated plasmids and performed protein purifications and binding assays; T.MK. and H.K. performed co-sedimentation assays; T.F. performed mass spectrometry analysis; L.K., M.Hi. and M.Ha performed immune inhibition assays. C.S. contributed and designed codon-optimized plasmids. M.T., R.Bu., M.A., T.B., F.K., K.S. and R.Be. wrote the manuscript, with comments from all authors.

## Data availability

Cryo-EM volumes and molecular models have been deposited at the Electron Microscopy Data Bank and Protein Data Bank with accession codes EMD-XXXX, EMD-XXXX, EMD-XXXX, EMD-XXXX, EMD-XXXX, EMD-XXXX, EMD-XXXX, EMD-XXXX, EMD-XXXX, EMD-XXXX, and PDB-YYYY, PDB-YYYY, PDB-YYYY, PDB-YYYY, PDB-YYYY.

## Supplementary Materials

Materials and methods

Figures S1-S5

Tables S1-S2

References (47-61)

