## Supplementary Materials for "Structural basis for translational shutdown and immune evasion by the Nsp1 protein of SARS-CoV-2"

### Materials and Methods

#### Cell lines, plasmids, viruses and transfections

HEK293T cells were purchased from American type culture collection (ATCC: #CRL-3216) and cultivated in Dulbecco's Modified Eagle Medium (DMEM, Gibco) supplemented with 10% (v/v) fetal bovine serum (FBS, Gibco), 100 U/ml penicillin (PAN-Biotech), 100 µg/ml streptomycin (PAN-Biotech) or 1x penicillin/streptomycin (Gibco), and 2 mM L-glutamine (PAN-Biotech) or 1x GlutaMAX (Gibco) (hereafter called DMEM+3).

The open reading frames for SARS-CoV-2 (SCoV-2) Nsp1and Nsp7 were ordered codon-optimized and V5-tagged from Twist Bioscience and the SARS-CoV (SCoV-1) and SCoV-2 Nsp1 open reading frames were in addition synthesized by GenArts (ThermoFisher Scientific). Nsp1-V5 and Nsp7-V5 were amplified by PCR (using Primer 1 (Nsp1 rev): CGA CGC GTC TAG CCG CCA TTC AGC TCG CGC, primer 2 (Nsp7 rev): CGA CGC GTC TAT TGC AGC GTG GCA CG, primer 3 (XbaI fwd) CGT CTA GAG CCA CCATG) and the single restriction sites XbaI/MluI as well as the Kozak sequence GCCACC were introduced. Afterwards PCR-fragments were subcloned into pCG or pCG-IRES-GFP vectors to yield pCGSARS-CoV-2 Nsp1-V5 IRES eGFP, pCGSARS-CoV2 Nsp7-V5 IRES eGFP, pCGSARS-CoV-2 Nsp1-V5, pCGSARS-CoV-2 -Nsp7-V5. pCR3-MeV-V-FLAG, pIFNb-FFLuc, pISRE-FFLuc, pCMV-GLuc were kind gifts from Karl-Klaus Conzelmann and described previously (*40*). The SCoV-1 Nsp1 and SCoV-2 Nsp1 open reading frames were PCR amplified with Primer 4 (BamHI Nsp1 SCoV-1) CGC GGA TCC ATG GAG AGC CTT GTT CTT GGT GTC AAC G, Primer 5 (Nsp1 STOP SCoV-1 XhoI) CCG CTC GAG TCA ACC TCC ATT GAG CTC ACG AGT GAG TTC and Primer 6 (BamHI Nsp1 SCoV-2) CGC GGA TCC ATG GAG AGC CTT GTC CCT GGT TTC AAC G, Primer 7 (Nsp1 STOP SCoV-2 XhoI) CCG CTC GAG TCA CCC TCC GTT AAG CTC ACG CAT GAG TTC and cloned into a modified pcDNA5/FRT/TO-3xFLAG-3C expression vector for N-terminal tagging. In order to generate *E.coli* expression plasmids Nsp1 from SCoV-1 and SCoV-2 were amplified with Primer 8 (NdeI Nsp1 SCoV-1) GGG AAT TCC ATA TGG AGA GCC TTG TTC TTG GTG TCA ACG, Primer 9 (Nsp1 STOP SCoV-1 BamHI) CGC GGA TCC TTA ACC TCC ATT GAG CTC ACG AGT GAG TTC and Primer 10 (NdeI Nsp1 SCoV-2) GGG AAT TCC ATA TGG AGA GCC TTG TCC CTG GTT TCA ACG, Primer 11 (Nsp1 STOP SCoV-2 BamHI) CGC GGA TCC TTA CCC TCC GTT AAG CTC ACG CAT GAG TTC and were cloned into the plasmid backbones pET-24d-(His)6-TEV and pET-24d-GST-TEV for N-terminal tagging. The K164A H165A mutations were introduced by overlap extension PCR using the following primer: Primer 12 (Nsp1 KH>AA SCoV-1_fwd) TGG AAC ACT GCC GCT GGC AGT GGT GCA CTC CGT GAA CTC A, primer 13 (Nsp1 KH>AA SCoV-1_rev) CTG CCA GCG GCA GTG TTC CAG TTT TGT TCA TAA TCT TCA ATG GG and Primer 14 (Nsp1 KH>AA SCoV-2_fwd) TGG AAC ACT GCA GCT AGC AGT GGT GTT ACC CGT GAA CTC ATG, Primer 15 (Nsp1 KH>AA SCoV-2_rev) CTG CTA GCT GCA GTG TTC CAG TTT TCT TGA AAA TCT TCA TAA GG.

Firefly luciferase (Fluc) mRNA diluted in 25 µl Opti-Mem (Gibco) was mixed with 1 µl Lipofectamin2000 (Invitrogen) diluted in 25 µl Opti-Mem, incubated for 5 min at room temperature and transferred on the cells. For all transfections of plasmid DNA, Polyethylenimine (PEI, 1 mg/ml in H2O, Sigma-Aldrich) or the TransIT-LT1 Transfection Reagent (Mirus) were used according to the manufacturers recommendations or described previously (*47*).

Sendai virus (SeV, Cantell strain) was kindly provided by Georg Kochs (Freiburg University) and Daniel Sauter (Ulm University) and used to stimulate innate immune activation via RIG-I 16 h post transfection. Recombinant IFN beta was purchased from R&D Systems (8499-IF) and used at a concentration of 1000 U/ml for stimulation either for 8 h (luciferase reporter gene assays) or for 24 h (ISG expression).

Whole-cell lysates

Whole-cell lysates were prepared by collecting cells in Phosphate-Buffered Saline (PBS, Gibco). The cell pellet (500 g, 4°C, 5 min) was lysed in transmembrane lysis buffer [50 mM HEPES pH 7.4, 150 mM NaCl, 1% Triton X-100, 5 mM ethylenediaminetetraacetic acid (EDTA)] by vortexing at maximum speed for 30 s. Cell debris were pelleted by centrifugation (20,000 g, 4°C, 20 min) and the cleared supernatants were stored for analysis at -20°C.

SDS-PAGE and immunoblotting

SDS-PAGE and immunoblotting was performed using standard techniques as previously described (*47*). The following antibodies were used throughout the study: anti-β-actin (1:10,000, AC-15, Invitrogen), anti-GAPDH (1:1000, # 607902, BioLegend), anti-V5 (1:1000, D3H8Q, Cell Signaling Technology), and anti-FLAG (1:1000 or 1:5000, M2, Sigma-Aldrich), IRDye Secondary Antibodies, Li-Cor (1:20,000 in 0.05% (w/v) casein, Thermo Scientific).

Legendplex ELISA

The Legendplex ELISA (Anti-virus Panel, Biolegend) was performed according to the manufacturer's instructions. In brief, the supernatants were incubated for 2 h at room temperature with the antibody-coated beads, followed by washing and incubation with the detection antibodies. After incubation with the staining reagent, the beads were analysed in a high-throughput sampler via flow cytometry (Canto II, BD Sciences). Absolute quantification was performed using a standard and the Biolegend Legendplex v8.0 software.

Luciferase assay

16 h after mRNA transfection or 24 h post transfection, the cells were lysed in 200 µl 1x Passive Lysis buffer (Promega). 25 µl of the lysate were transferred into a white 96-well plate, 25 µl of firefly substrate (Promega) were added and the luminescence was quantified as relative light units (Orion microplate luminator). To measure Gaussia luciferase activity for normalisation, 25 µl of the supernatants were transferred into a white 96-well plate. Coelenterazine Substrate (pjk), diluted 1:120 in PBS, was added to the samples and luminescence quantified (Orion microplate luminator).

qRT-PCR

Total RNA was extracted from cells using the Quick-RNA Microprep Kit (Zymo Research) according to the manufacturer's instructions. Reverse transcription and qRT–PCR were performed in one step using 2 µl (~400 ng) of the purified RNA samples as templates (SuperScript III Platinum Kit, Invitrogen) on a StepOnePlus Real-Time PCR System (Applied Biosystems) according to the manufacturer's instructions. TaqMan probes for each individual gene were acquired as premixed TaqMan Gene Expression Assays (Thermo Fisher Scientific) and added to the reaction. Expression level for each target gene was calculated by normalizing against GAPDH using the ΔΔCT method and represented relative to the values for mock-transfected cells, which were set to 1.

*In vitro* transcription (IVT)

For IVT of the luciferase reporter, the HiScribe T7 Quick High Yield RNA Synthesis Kit und Cap analogue (both NEB) were used. Capped RNA synthesis, DNAse treatment und LiCl precipitation were performed according to the supplier's protocol. As a template, the included linearized Firefly Luciferase template DNA was used. For IVT of the hCMV stalling mRNA (*in vitro* translation assay) the mMESAGE mMACHINE T7 Transcription Kit was used according to the manufacturer's protocol. Template for hCMV stalling mRNA was PCR amplified from plasmid pGEM4-CD4-CMV. The size of the all transcripts was analysed by agarose gel electrophoresis, stained using ethidium bromide (AppliChem GmbH) and visualized in a Bio-Rad Gel Doc XR+.

Quantification of autophagy

Transiently transfected HEK293T cells stably expressing GFP-LC3B were harvested in PBS and treated for 20 min at 4°C with PBS containing 0.05% Saponin to wash out non-membrane bound GFP-LC3B. Cells were subsequently fixed in 4% Paraformaldehyde (Santa Cruz) and fluorescence intensity quantified via flow cytometry (BD Canto II). The GFP-LC3B mean fluorescence intensity of the control was subtracted.

*In vitro* translation assay

HeLa S3 translation extract was prepared as described before (*48*), except that cells were treated with 200 nM integrated stress response inhibitor (ISRIB) 1 h prior to harvesting to ensure cap-dependent translation initiation (*49*). For the reaction, 12 µl HeLa translation reaction mix with 50% (v/v) extract was adjusted to 2.75 mM $Mg(OAc)_2$, 0.42 mM $MgCl_2$, 75 mM KOAc, 37.5 mM KCl, 42 mM NaCl, 2 mM DTT, 1.56 mM GTP, 0.25 mM ATP, 1.6 mM creatine phosphate, 0.45 mg/mL creatine kinase, 50 µg/mL yeast tRNA, 0.4 mM spermidine, 0.12 mM complete amino acid mixture (Promega) and 0.8 U/µl RNase inhibitor (Invitrogen).

Crude rabbit reticulocyte lysate (RRL, Green Hectares) was first treated with micrococcal nuclease, then supplemented with Hemin (*50*) and frozen in aliquots. The final 25 µl translation reaction mixture with 70% (v/v) RRL was adjusted to 50 mM NaCl, 0.5 mM $MgCl_2$, 0.3 mM complete amino acid mixture (Promega), 90 mM creatine phosphate, 0.1 mg/mL yeast tRNA and 7.3 U/µl SUPERase·In RNase inhibitor (Invitrogen).

HeLa and RRL translation reactions were preincubated with purified N-terminal $(His)_6$-TEV tagged Nsp1 or Nsp1-mt from SARS-CoV-1 and -2 for 30 min on ice with final concentrations of 6.25 µM (HeLa) and 7.5 µM (RRL). Subsequently, the reactions were initiated by addition of 1 µg of mRNA encoding the gp48 uORF2 peptide that leads to stalling of eukaryotic ribosomes (fig. S1B). Both *in vitro* translation reactions were incubated for 20 min at 30°C. Subsequently, ribosomes were isolated from the RRL reaction by pelleting through a sucrose cushion [50 mM HEPES-KOH pH 7.5, 10 mM $Mg(OAc)_2$, 200 mM KOAc, 1 mM DTT, 0.5 M sucrose] in a TLA100 rotor for 1 h at 434,513 g and 4°C. The ribosome pellets were resuspended in reducing sample buffer [50 mM Tris-HCl pH 6.8, 2% SDS, 0.005% bromophenol blue, 10% glycerol, 100 mM DTT] and separated using SDS-PAGE. Samples from HeLa translation reactions were loaded directly, without pelleting. Peptidyl-tRNA was detected by immunoblotting with primary antibodies against the V5-tag and secondary HRP-coupled antibodies.

Polyribosome gradient analysis

HEK293T were cultured to 40% confluency and transiently transfected with the pcDNA/FRT/TO-3xFLAG-3C-Nsp1 plasmids using PEI. After 22 h cells from one 15 cm culture dish (~$1.8x10^7$) were treated with 100 µg/ml cycloheximide (CHX) for 15 min, then washed with 5 ml ice-cold PBS containing 100 µg/ml CHX and scraped off the culture dish. Cells were harvested by centrifugation at 120 x g for 10 min. The cells were washed once with 1 ml PBS/CHX and pelleted by centrifugation at 300 x g for 5 min. The cell pellet was resuspended in 500 ml lysis buffer [5 mM Tris-HCl, pH 7.5, 1.5 mM KCl, 2.5 mM $MgCl_2$, complete EDTA-free protease inhibitor cocktail (Roche), 100 µg/ml CHX, 2 mM DTT, 200 U/ml SUPERase·In RNase Inhibitor (Invitrogen), 0.5% v/w Triton X-100, and 0.5% v/w sodium deoxycholate], incubated for 1 minute and cleared by centrifugation at 15000 x g for 5 min. The nucleic acid concentration in the cleared samples was determined by measurement of the absorption at 260 nm and an equivalent amount of each sample was separated on a 10%-50% sucrose gradient [20 mM HEPES-KOH, pH 7.5, 100 mM KOAc, 5 mM $MgCl_2$, 1 mM DTT, 10 U/ml RNase inhibitor, protease inhibitor cocktail] at 202048 x g for 2.5 h using a SW 40 Ti rotor (Beckman Coulter). Gradients were fractionated using a Biocomp piston gradient fractionator and $A_{260}$ was observed using a Biocomp Triax flow cell. Proteins in the collected fractions were precipitated using trichloroacetic acid and separated on a 15% SDS-PAGE gel, then blotted onto a PVDF membrane (0.45 µm pore size, Immobilon-P, Merck). Detection of 3xFLAG-3C-Nsp1 on the membranes was performed using an M2 anti-FLAG HRP antibody (Sigma, A8592) at 1/1000 working dilution according to the manufacturer's protocol. Fractions from one gradient were blotted on two separate membranes, decorated with AB at the same time, and developed using the same imaging parameters. Images were then combined, and band intensities equalized based on samples loaded on both membranes.

Expression and purification from *E. coli*

The constructs were transformed and expressed in *E. coli* BL21(DE3) cells. Cells were grown at 37°C in LB medium supplemented with kanamycin (50 µg/ml). When the cultures reached an $OD_{600}$ of about 0.5-0.7, cells were shifted to 18°C and expression was induced after 30 min by the addition IPTG to a final concentration of 0.25 mM. Cells were grown for 16 h, harvested by centrifugation and stored at -80°C. For purification of the GST-TEV tagged Nsp1 constructs cell pellets were resuspended in lysis buffer [20 mM HEPES-NaOH pH 7.5, 500 mM NaCl, 5 mM $MgCl_2$] and lysed with a M-110L Microfluidizer (Microfluidics). The lysate was cleared by centrifugation (43,200 x g, 30 min, 4°C) and the supernatant was incubated with Glutathione Sepharose 4 Fast Flow beads (GE Healthcare) for 1 h at 4°C. The supernatant was removed and beads were washed two times with 40 ml lysis buffer. Samples were eluted with lysis buffer supplemented with 25 mM reduced L-gluthathione (Sigma-Aldrich). Cells expressing the different $(His)_6$-TEV-Nsp1 and the Nsp1-AviTag-$(His)_6$ construct were resuspended and lysed in lysis buffer 2 [20 mM HEPES-NaOH pH 8.0, 500 mM NaCl, 5 mM $MgCl_2$, 10 mM imidazole] as described above. Ni-NTA agarose (Qiagen) was added to the cleared lysate and incubated for 1 h at 4°C. Beads were washed once with 40 ml lysis buffer 2 and once with 30 ml lysis buffer 2 containing 20 mM imidazole. Samples were eluted with buffer containing: 20 mM HEPES-NaOH pH 7.5, 500 mM NaCl, 5 mM $MgCl_2$ and 300 mM imidazole. All samples were further purified by size-exclusion chromatography in buffer containing: 20 mM HEPES-NaOH pH 7.5, 500 mM NaCl, 5 mM $MgCl_2$. The $(His)_6$-TEV-Nsp1 and the Nsp1-AviTag-$(His)_6$ proteins were purified through a Superdex 75 10/300 GL column (GE Healthcare) and for the GST-TEV-Nsp1 samples a Superdex 200 Increase 10/300 GL (GE Healthcare) column was used. Fractions containing Nsp1 were pooled, concentrated with a Amicon Ultra centrifugal filter (Millipore), flash frozen in liquid nitrogen and stored at -80°C.

In vitro binding assay

Human 40S and 60S ribosomal subunits were prepared as previously described (*24*). The binding assay was performed in binding buffer [20 mM HEPES-KOH pH 7.6, 150 mM KOAc, 2.5 mM $MgCl_2$, 0.01% NP-40 and 2 mM DTT]. The GST-TEV tagged Nsp1 bait proteins (60 pmol) were incubated with purified 40S or 60S ribosomal subunits (see fig. S1A) in a total volume of 300 µl for 45 min at 4°C. The samples were transferred to 1 ml Mobicols (MoBiTec) containing 30 µl Glutathione Sepharose 4 Fast Flow (GE Healthcare) and incubated for an hour at 4°C. The unbound material was removed by centrifugation in a pre-cooled table top centrifuge and the beads were washed 3x times with binding buffer (1x 800 µl, 2x 500 µl). The beads were incubated for 1 h at 4°C with binding buffer supplemented with 25 mM reduced L-glutathione (Sigma-Aldrich) and the samples were eluted by centrifugation. Samples were analyzed on a 12% polyacrylamide gel (NuPAGE, Invitrogen) and stained with Coomassie.

Affinity purification of Nsp1 from HEK293T cells

HEK293T cells transiently transfected with the pcDNA5/FRT/TO-3xFLAG-3C-Nsp1 (SCoV-2) construct were lysed in purification buffer [20 mM Hepes-KOH pH 7.5, 150 mM KOAc, 5 mM $MgCl_2$, 1 mM DTT, 0.5 mM NaF, 0.1 mM $Na_2V_3O_4$] supplemented with 5% glycerol and 0.5% NP-40. The lysate was sonicated 4 times for 10 s with 30 s on ice in between (Branson Sonifier 250). The lysate was cleared by centrifugation for 15 min at 2,960 g and 25 min at 36,500 g and the supernatant was incubated with ANTI-FLAG M2 agarose beads (Sigma-Aldrich) on a rotating wheel for 120 min at 4°C. The FLAG beads were washed twice with 10 ml purification buffer supplemented with 0.01% NP-40 and once with purification buffer containing 0.05% Nikkol. The FLAG beads were transferred to a 1 ml Mobicol (MoBiTec) and washed with 5 ml buffer + 0.05% Nikkol. The beads were incubated with buffer containing 20 mM HEPES-KOH pH 7.5, 150 mM KOAc, 5 mM $MgCl_2$, 0.05% Nikkol and 40 µg 3C protease for 1 h at 4°C and the eluted samples were collected by centrifugation. Eluates were used for Cryo-EM analysis and analyzed on a 4-12% Bis-Tris polyacrylamide gel (NuPAGE, Invitrogen).

Mass spectrometry

The gel bands were de-stained using 50% acetonitrile in 50 mM $NH_4HCO_3$. For protein reduction, 45 mM dithioerythritol (DTE) in 50 mM NH4$HCO_3$ was added to the gel pieces and incubated for 30 min at 55°C. Carbamidomethylation of cysteines was done twice by incubation at RT in the dark for 15 min in 100 mM iodoacetamide/50 mM $NH_4HCO_3$. Gel slices were minced and digestion was performed overnight at 37°C using 70 ng porcine trypsin (Promega, Fitchburg, WI, USA). Tryptic peptides were extracted using 70% ACN. Prior to liquid chromatography, the samples were dried using a SpeedVac vacuum concentrator. Chromatography was done with an Ultimate 3000 nano-LC system (Thermo Fisher Scientific) using an Acclaim PepMap 100 trap column (nanoViper C18, 2 cm length, 100 µM ID, Thermo Scientific) and an EasySpray separation column (PepMap RSLC C18, 50 cm length, 75 µm ID, Thermo Fisher Scientific). For peptide separation a flow rate of 250 nl/min and 0.1% formic acid as solvent A was used. The method consisted on gradients from 3% to 25% solvent B (0.1% formic acid in acetonitrile) in 30 min and from 25% to 40% B in 5 min. Data dependent mass spectrometry was performed on a Q Exactive

HF-X mass spectrometer (Thermo Fisher Scientific) using cycles of one full scan (350 to 1600 m/z) at 60k resolution and up to 12 data-dependent MS/MS scans at 15k resolution. Spectra were searched using MASCOT V2.4 (Matrix Science Ltd, London, UK) and the human subset of the UniProt database. Common contaminants like keratins were removed and the results were filtered for an FDR < 1%. Exclusively protein identifications with at least two individual peptides were considered.

Electron microscopy and image processing

3.5 µl of sample solution were applied to holey carbon support grids (R3/3 with 2 nm continuous carbon support, Quantifoil), which had been glow discharged at $2.1x10^{-1}$ mbar for 20 s. Grids were incubated for 45 s at 4°C and subsequently plunge frozen in liquid ethane using a Vitrobot Mark IV (FEI Company). 11,270 (dataset 1, reconstituted complex) and 6,610 (dataset 2, native complex) movies were recorded on a Titan Krios at 300 kV using a K2 Summit direct electron detector with a nominal pixel size of 1.059 Å and a defocus range from 0.5 – 3.0 µm at low-dose conditions. For each movie, 40 frames with approximately 1.12 $e^{-}$ $Å^{-2}$ exposure were gain corrected and aligned using MotionCor2 (*51*). Contrast-transfer function (CTF) parameters of the summed micrographs were estimated with Gctf (*52*) and CTFFIND4 (*53*), before micrographs were manually screened for quality. 1,690,969 (dataset 1) and 701,423 (dataset 2) particles from 11,270 (dataset 1) and 5,799 (dataset 2) micrographs were then picked using Gautomatch (dataset 1), or the Relion 3.1 AutoPick function (dataset 2) (*54, 55*). The particles from dataset 1 displayed severe orientation bias and were further processed using cryoSPARC (*56*). The particles from dataset 2 were further processed in Relion 3.1. All particles were subjected to extensive 2D and 3D classification in cryoSPARC (dataset 1) and Relion 3.1 (dataset 2) (fig. S2, C and D). Particles were then subjected to CTF parameter refinement in cryoSPARC (dataset 1) and Relion 3.1 (dataset 2), before final reconstructions were prepared. In case of the reconstituted Nsp1-40S complex (dataset 1), local refinements were performed in cryoSPARC using masks on the body and head of the 40S. The focused refined maps were filtered according to local resolution and sharpened using a B-factor of -80, before they were combined to a composite map using Phenix (*57*). Furthermore, both 43S PIC volumes (dataset 2) were subjected to multi-body refinement in Relion 3.1 using a masks on the 40S body, head and eIF3, as well as eIF2-tRNA in case of 43S PIC state 2. Finally, the local resolution of each reconstituted volume was estimated using cryoSPARC or Relion 3.1.

Statistical calculations

Unpaired student's t-test (Welch correction) were used to calculate significances for luciferase assays, qPCRs and ELISA. Not significant values are indicated as ns; Significant samples denoted as *,p<0.01;**, p<0.001; ***, p<0.0001.

Model building and refinement.

The molecular model of Nsp1-40S was manually built in Coot (*58, 59*) using PDB-6G5H as an initial model of the small ribosomal subunit. All proteins and ribosomal RNA were checked and refined in Coot, before the C-terminal helices of Nsp1 were built *de novo*. The model was subsequently real-space refined in Phenix and the surface potential of Nsp1-C determined using DelPhi (*60*). Models of all Nsp1-80S ribosomal complexes were based on the structure of CCDC124-bound 80S ((*30*), unpublished). C-terminal domains of CCDC124 and LYAR were built de novo in Coot. Models for ABCE1 and eRF1 were based on PDB-5LZV and rigid-body fit into the density before manually adjusting domain 3 of eRF1 in Coot. Models for the ternary eEF1A-tRNA-GTP complex were based on PDB-5LZS and rigid-body fitted into density. The sequence of pre-accommodated tRNA was manually changed to leucyl-tRNA and the extended variable loop built. All models were then real-space refined once in Phenix.

Cryo-EM densities and molecular models were visualized using ChimeraX (*61*).

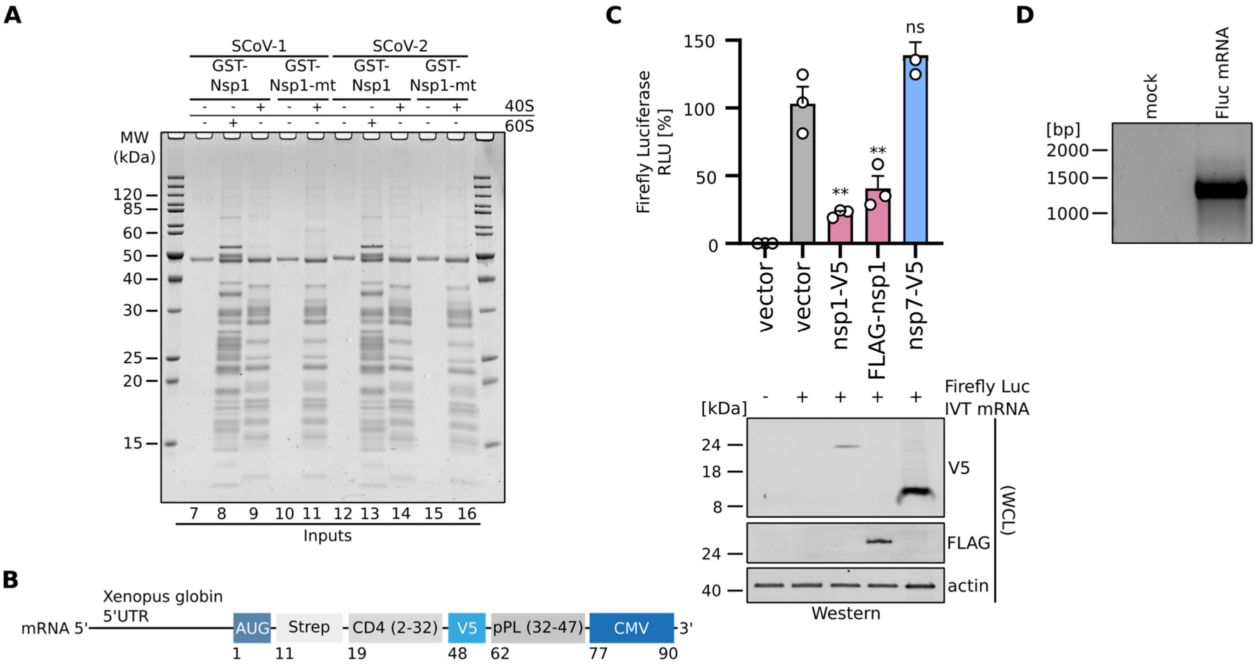


**Fig. S1. Characterization of Nsp1 from SARS-CoV and SARS-CoV-2.**
(**A**) Input samples of the *in vitro* binding assay described in Fig. 2, B. The samples were analyzed by SDS-PAGE and Coomassie staining. SCoV-1, SARS-CoV. SCoV-2, SARS-CoV-2. (**B**) Organization of the reporter mRNA construct used in the *in vitro* translation assay in Fig. 1, D: mRNA coding for an N-terminal Strep-tag, amino acids 2-32 of CD4, a V5-tag, amino acids 32-47 of preprolactin (pPL) and a sequence coding for the arrest peptide of gp48 uORF2 from cytomegalovirus (CMV). (**C**) Luciferase reporter gene assay of HEK293T cells transiently transfected with indicated plasmids (empty vector, Nsp1-V5, FLAG-Nsp1-FLAG, Nsp7-V5) and *in vitro* transcribed Firefly luciferase mRNA. Bars represent the mean of n=3±SEM. RLU, relative light units, normalized to empty vector set to 100%. (top panel). Immunoblot of whole cell lysates of the reporter gene assay stained with anti-V5, anti-FLAG and anti-actin antibodies (bottom panel). (**D**) Agarose gel of the *in vitro* transcribed Firefly luciferase mRNA used in Fig. 1E.

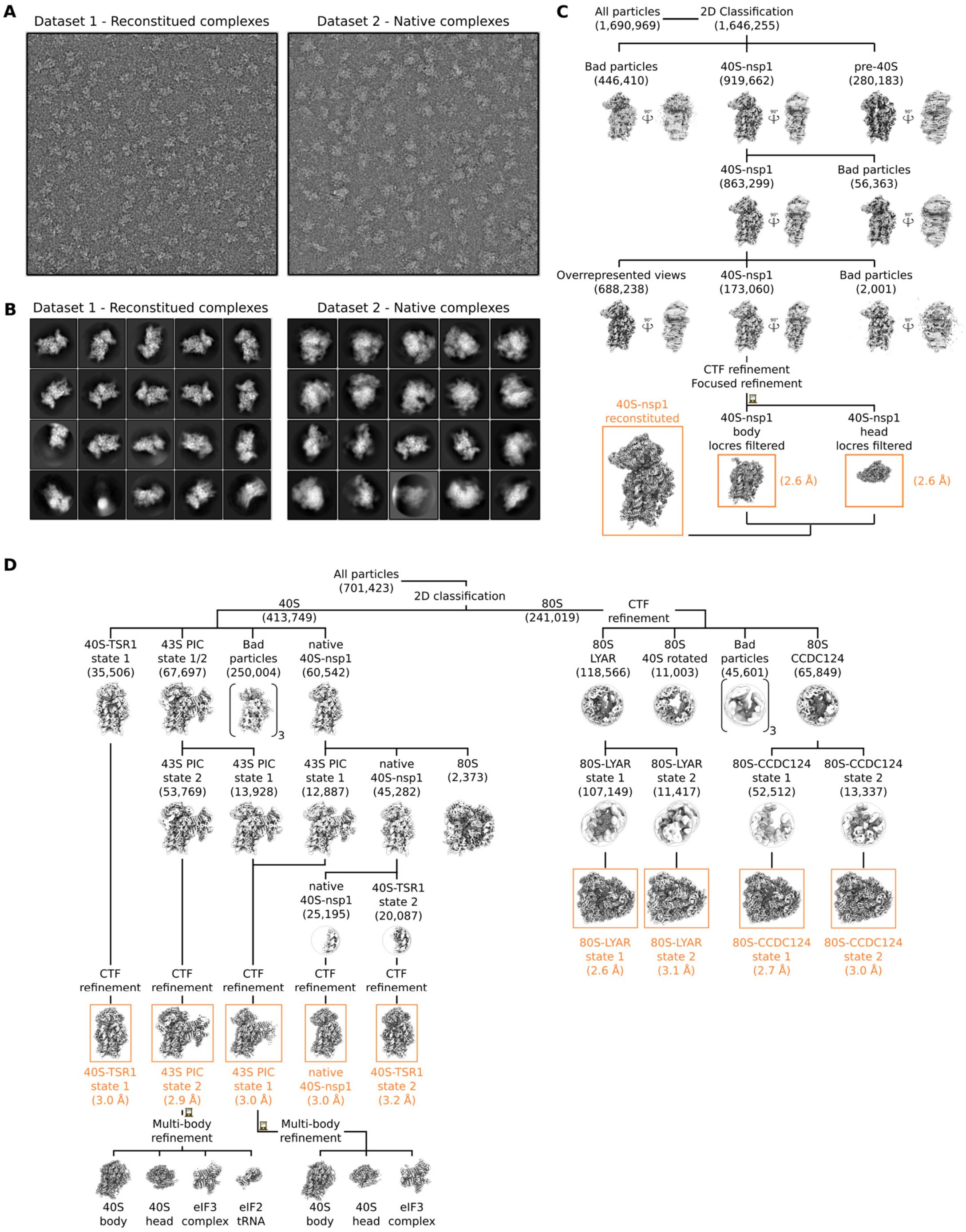


**Fig. S2. Cryo-electron microscopy analysis.**
(**A**) Representative electron micrographs from both reconstituted and native datasets, respectively, displayed with inverted contrast and after low-pass filtering at 20 Å. (**B**) Selected image averages after particle image 2D classification. (**C** and **D**) Cryo-EM data processing scheme for the reconstituted Nsp1-40S dataset (**C**) and affinity purified ribosomal complexes (**D**). Respective image numbers are provided in brackets and final volumes and their resolution marked in orange.

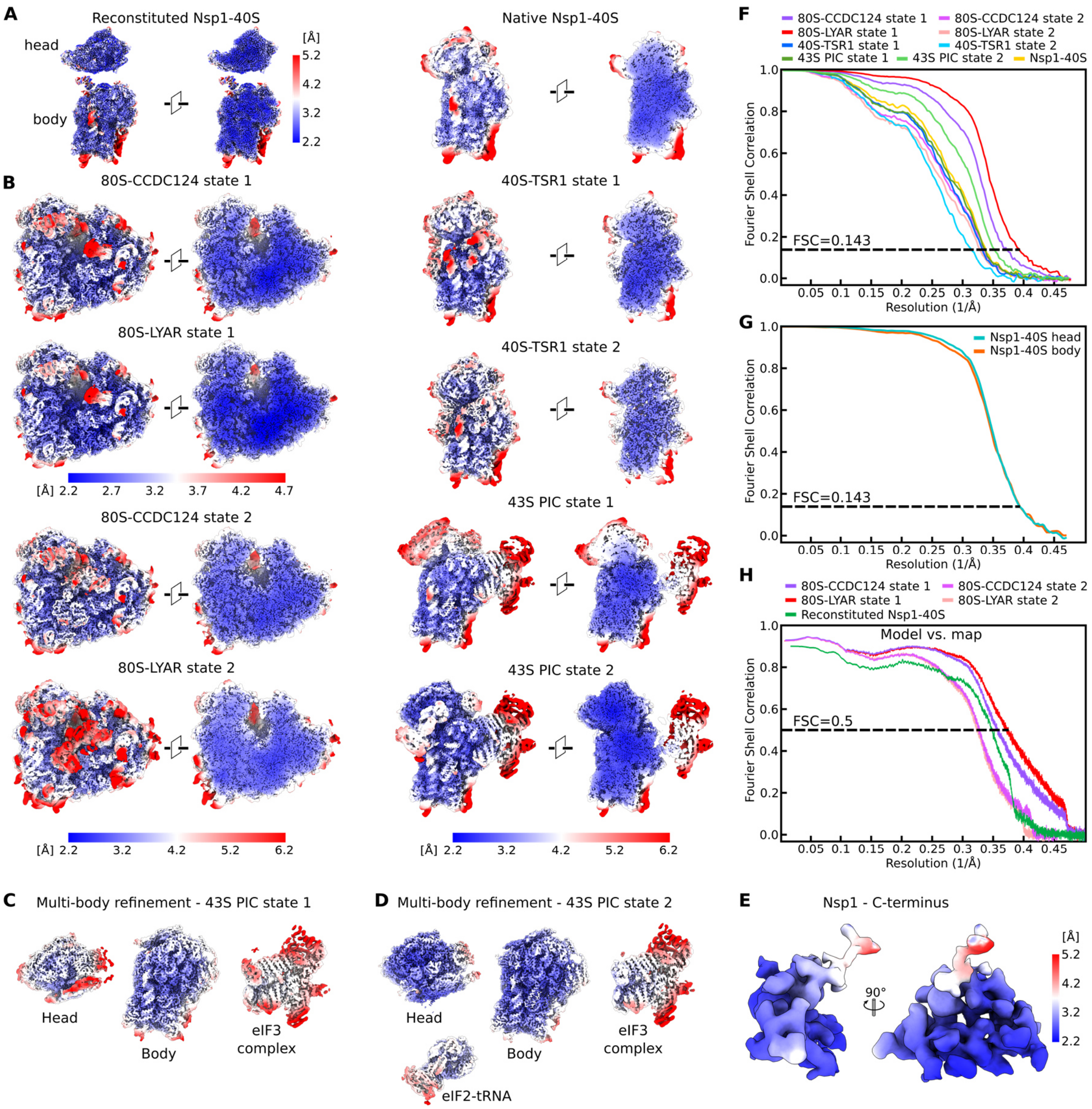


**Fig. S3. Local resolution and model statistics.**
(**A** to **E**) Final cryo-EM maps filtered and colored according to local resolution as estimated by cryoSPARC (**A** and **E**) and Relion (**B** to **D**). Reconstituted Nsp1-40S (**A**) and both 43S PIC states (**C** and **D**) were locally or multi-body refined in cryoSPARC and Relion, respectively, and their respective subvolumes are shown. Colors refer to local resolution averages as shown by the nearest color key to the right (**A** and **C** to **E**) or below each image (**B**). (**F** and **G**) Fourier shell correlation (FSC) curves of all native Nsp1-ribosomal complexes (**F**) and locally refined maps of reconstituted Nsp1-40S (**G**) as estimated by Relion. Threshold for final resolution estimation according to the 'gold-standard' set at FSC=0.143. (**H**) FSC curve of the final models against their respective cryo-EM maps as provided by Phenix.

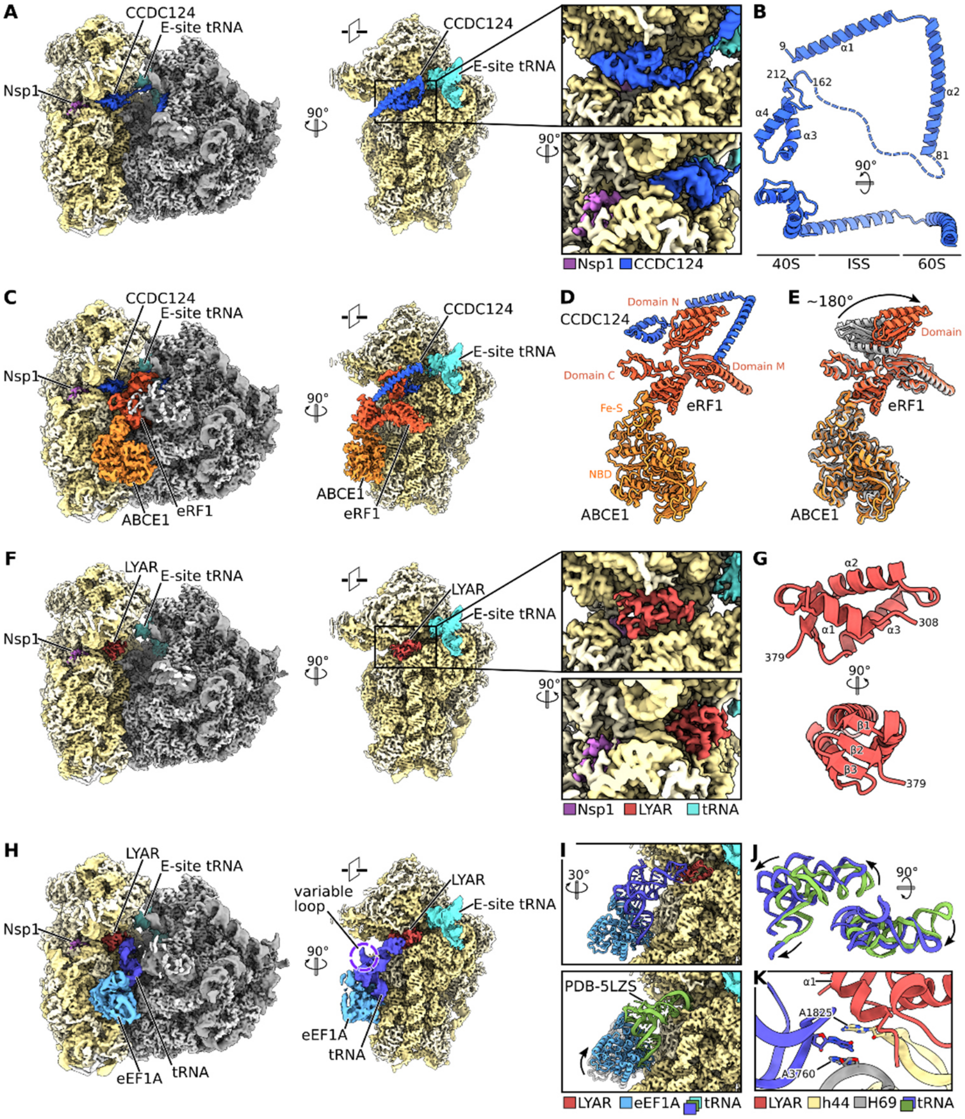


**Fig. S4. Nsp1-bound 80S ribosomal complexes display unusual factor compositions.**
(**A**) Cryo-EM map of Nsp1-80S bound by CCDC124 with its C-terminal domain positioned at the A-site. (**B**) Molecular model of CCDC124. CCDC124 forms two long alpha-helices (α1 and α2), which span the intersubunit side (ISS) and bind to the 60S as previously described (*30*). The newly observed C-terminal domain comprises two alpha-helices (α3 and α4) and is linked to α2 by a long flexible linker (dashed line). (**C**) Cryo-EM volume of CCDC124 bound Nsp1-80S with splitting and translation termination factors ABCE1 and eRF1. (**D**) Molecular models of ABCE1, eRF1 and CCDC124 with their domains labeled show the central position of eRF1 domain N encompassed by CCDC124. (**E**) Superposition of ABCE1-eRF1 from the Nsp1-80S volume (red and orange) and PDB-3JAH (gray) highlighting the rotated position of eRF1 domain N. Domain N, which is usually positioned within the A-site decoding the stop codon, is displaced by the C-terminal fold of CCDC124. (**F**) Cryo-EM volume of a Nsp1-80S complex, containing LYAR. LYAR occupies the mRNA channel at the A-site with its C-terminal fold. (**G**) Three alpha-helices (α1 – α3) and a three-stranded beta-sheet (β1 – β3) form the observed part of LYAR in a α1-β1-α2-α3-β2-β3 sequence. (**H**) Cryo-EM volume of a Nsp1-80S ribosome with LYAR bound by the ternary

eEF1A-tRNA-GTP complex. The large variable loop of the pre-accommodated type II tRNA is marked. (**I**) Molecular model of eEF1A, tRNA and LYAR bound to the 40S subunit volume shows the anticodon loop displaced by LYAR (top). Model of a canonical elongation complex (PDB-5LZS) docked into the cryo-EM map shows a different position of eEF1A and tRNA (bottom). (**J**) Superposition of the tRNA in the Nsp1-80S state and PDB-5LZS highlights the repositioned tRNA with a stretched decoding loop. (**K**) Molecular interactions between LYAR, tRNA and rRNA helices h44 and H69. Base stacking occurs between A3760 of the 28S rRNA, A1825 of the 18S rRNA and a base of the anticodon loop of the bound tRNA. A model of a leucyl-tRNA was used, however, local resolution prevented unambiguous identification of the bound tRNA.

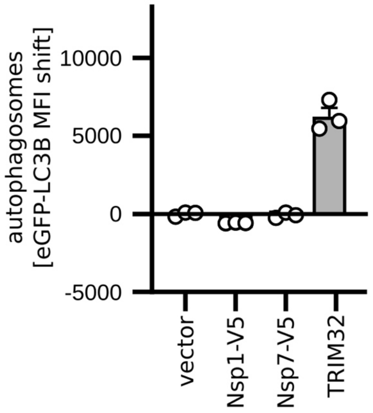


**Fig. S5. Modulation of autophagy by SARS-CoV-2 proteins**

Autophagy levels in HEK293T cells stably expressing GFP-LC3B and transiently transfected with indicated plasmids (empty vector, nsp1-V5, nsp7-V5 and TRIM32-FLAG). Autophagosomes were quantified by flow cytometry as mean fluorescence intensity of GFP-LC3B-positive vesicles saponin-permeabilised cells. Bars represent the mean of n=3±SEM. The MFI of the vector control was set to 0.

**Table S1. Data collection, refinement and model statistics**

| | Nsp1-40S reconstituted (EMDB-XXX) (PDB-XXX) | 80S-CCDC124 state 1 (EMDB-XXX) (PDB-XXX) | 80S-CCDC124 state 2 (EMDB-XXX) (PDB-XXX) | 80S-LYAR state 1 (EMDB-XXX) (PDB-XXX) | 80S-LYAR state 2 (EMDB-XXX) (PDB-XXX) |
|---|---|---|---|---|---|
| **Data collection and processing** | | | | | |
| Camera | Gatan K2 Summit | Gatan K2 Summit | Gatan K2 Summit | Gatan K2 Summit | Gatan K2 Summit |
| Magnification | 130,000 | 130,000 | 130,000 | 130,000 | 130,000 |
| Voltage (kV) | 300 | 300 | 300 | 300 | 300 |
| Electron exposure (e-Å-²) | 44.8 | 44.8 | 44.8 | 44.8 | 44.8 |
| Defocus range (µm) | 0.5 - 3.0 | 0.5-3.0 | 0.5-3.0 | 0.5-3.0 | 0.5-3.0 |
| Pixel size (Å) | 1.059 | 1.059 | 1.059 | 1.059 | 1.059 |
| Symmetry imposed | C1 | C1 | C1 | C1 | C1 |
| Micrographs collected (no.) | 11,270 | 6610 | 6610 | 6610 | 6610 |
| Initial particle images (no.) | 1,690,969 | 701,423 | 701,423 | 701,423 | 701,423 |
| Final particle images (no.) | 173,060 | 52,512 | 13,337 | 107,149 | 11,417 |
| Map resolution (Å) | 2.6 | 2.7 | 3.0 | 2.6 | 3.1 |
| FSC threshold | 0.143 | 0.143 | 0.143 | 0.143 | 0.143 |
| **Refinement** | | | | | |
| Initial model used (PDB) | 6G5H | | | | |
| Model resolution (Å) | 2.7 | 2.8 | 3.0 | 2.7 | 3.1 |
| FSC threshold | 0.5 | 0.5 | 0.5 | 0.5 | 0.5 |
| Map sharpening Bfactor (Å2) | -80 | -20 | -20 | -20 | -20 |
| Model composition | | | | | |
| Non-hydrogen atoms | 74,562 | 225,552 | 233,406 | 225,112 | 230,342 |
| Protein residues | 4,887 | 12,547 | 13,537 | 12,495 | 1,935 |
| Nucleotide residues | 1,665 | 5,864 | 5,864 | 5,864 | 5,951 |
| Ligands | 3 | 264 | 267 | 264 | 264 |
| R.m.s deviations | | | | | |
| Bond lengths (Å) | 0.008 | 0.008 | 0.007 | 0.006 | 0.008 |
| Bond angles (°) | 1.083 | 0.924 | 1.037 | 0.864 | 0.939 |
| Validation | | | | | |
| Molprobity score | 1.44 | 2.38 | 2.47 | 2.32 | 2.51 |
| Clash score | 3.70 | 5.41 | 6.35 | 4.96 | 6.54 |
| Poor rotamers (%) | 0.40 | 6.39 | 7.03 | 5.82 | 7.20 |
| Ramachandran plot | | | | | |
| Favored (%) | 95.05 | 92.34 | 92.14 | 92.32 | 91.70 |
| Allowed (%) | 3.82 | 7.42 | 7.63 | 7.45 | 8.07 |
| Disallowed (%) | 0.23 | 0.24 | 0.22 | .024 | 0.24 |
| Map vs. Model CC (mask) | 0.82 | 0.86 | 0.82 | 0.86 | 0.82 |
| EMRinger score ($Nsp1^{148-180}$) | 5.44 | | | | |

**Table S2. Data collection statistics.**

| | Native 40S-Nsp1 (EMDB-XXX) | 40S-TSR1 state 1 (EMDB-XXX) | 40S-TSR1 state 2 (EMDB-XXX) | 43S PIC state 1 (EMDB-XXX) | 43S PIC state 2 (EMDB-XXX) |
|---|---|---|---|---|---|
| **Data collection and processing** | | | | | |
| Camera | Gatan K2 Summit | Gatan K2 Summit | Gatan K2 Summit | Gatan K2 Summit | Gatan K2 Summit |
| Magnification | 130,000 | 130,000 | 130,000 | 130,000 | 130,000 |
| Voltage (kV) | 300 | 300 | 300 | 300 | 300 |
| Electron exposure ($e^{-}Å^{-2}$) | 44.8 | 44.8 | 44.8 | 44.8 | 44.8 |
| Defocus range (μm) | 0.5-3.0 | 0.5-3.0 | 0.5-3.0 | 0.5-3.0 | 0.5-3.0 |
| Pixel size (Å) | 1.059 | 1.059 | 1.059 | 1.059 | 1.059 |
| Symmetry imposed | C1 | C1 | C1 | C1 | C1 |
| Micrographs collected (no.) | 6610 | 6610 | 6610 | 6610 | 6610 |
| Initial particle images (no.) | 701,423 | 701,423 | 701,423 | 701,423 | 701,423 |
| Final particle images (no.) | 25,195 | 35,506 | 20,087 | 13,928 | 53,769 |
| Map resolution (Å) | 3.0 | 3.0 | 3.2 | 3.0 | 2.9 |
| FSC threshold | 0.143 | 0.143 | 0.143 | 0.143 | 0.143 |

**Data S1. Mass spectrometry of SARS-CoV-2 Nsp1 purified from HEK293T**

Related to Fig. 2E
